# Steric Regulation of Tandem Calponin Homology Domain Actin-Binding Affinity

**DOI:** 10.1101/598359

**Authors:** Andrew R Harris, Brian Belardi, Pamela Jreij, Kathy Wei, Hengameh Shams, Andreas Bausch, Daniel A Fletcher

## Abstract

Tandem calponin homology (CH1-CH2) domains are common actin-binding domains in proteins that interact with and organize the actin cytoskeleton. Despite regions of high sequence similarity, CH1-CH2 domains can have remarkably different actin-binding properties, with disease-associated point mutants known to increase as well as decrease affinity for f-actin. To investigate features that affect CH1-CH2 affinity for f-actin in cells and in vitro, we perturbed the utrophin actin-binding domain by making point mutations at the CH1-CH2 interface, replacing the linker domain, and adding a PEG polymer to CH2. Consistent with a previous model describing CH2 as a steric negative regulator of actin binding, we find that utrophin CH1-CH2 affinity is both increased and decreased by modifications that change the effective ‘openness’ of CH1 and CH2 in solution. We also identified interface mutations that caused a large increase in affinity without changing solution ‘openness’, suggesting additional influences on affinity. Interestingly, we also observe non-uniform sub-cellular localization of utrophin CH1-CH2 that depends on the N-terminal flanking region but not on bulk affinity. These observations provide new insights into how small sequence changes, such as those found in diseases, can affect CH1-CH2 binding properties.

## INTRODUCTION

Actin filaments are organized into diverse cytoskeletal structures by a wide range of actin-binding proteins (Harris *et al*, 2018; Michelot & Drubin, 2011). Tandem calponin homology (CH1-CH2) domains are common actin-binding motifs found in diverse proteins, including the actin crosslinkers, α-actinin and filamin, as well as the membrane-actin linkers utrophin (utrn) and dystrophin (Korenbaum & Rivero, 2002; Bañuelos *et al*, 1998). Despite a conserved structural fold (Gimona *et al*, 2002) and regions of high sequence conservation (~20% identity and ~30% conservation across the tandem domain, Fig 1A), different CH1-CH2 domains bind to filamentous actin (f-actin) with affinities that can vary by an order of magnitude between closely-related proteins. For example, the affinity of α-actinin-1’s actin-binding domain (ABD) has a K_d_ = 4µM (Winder *et al*, 1995), while that of α-actinin-4’s ABD is >50µM (Lee *et al*, 2008). Similarly, filamin A’s ABD has a K_d_ = 47µM (Ruskamo & Ylänne, 2009) compared to 7µM for filamin B’s ABD (Sawyer *et al*, 2009), and utrophin’s ABD has a K_d_ = 19µM compared to 44µM for the ABD of its muscle homologue dystrophin (Winder *et al*, 1995).

**Figure 1:**
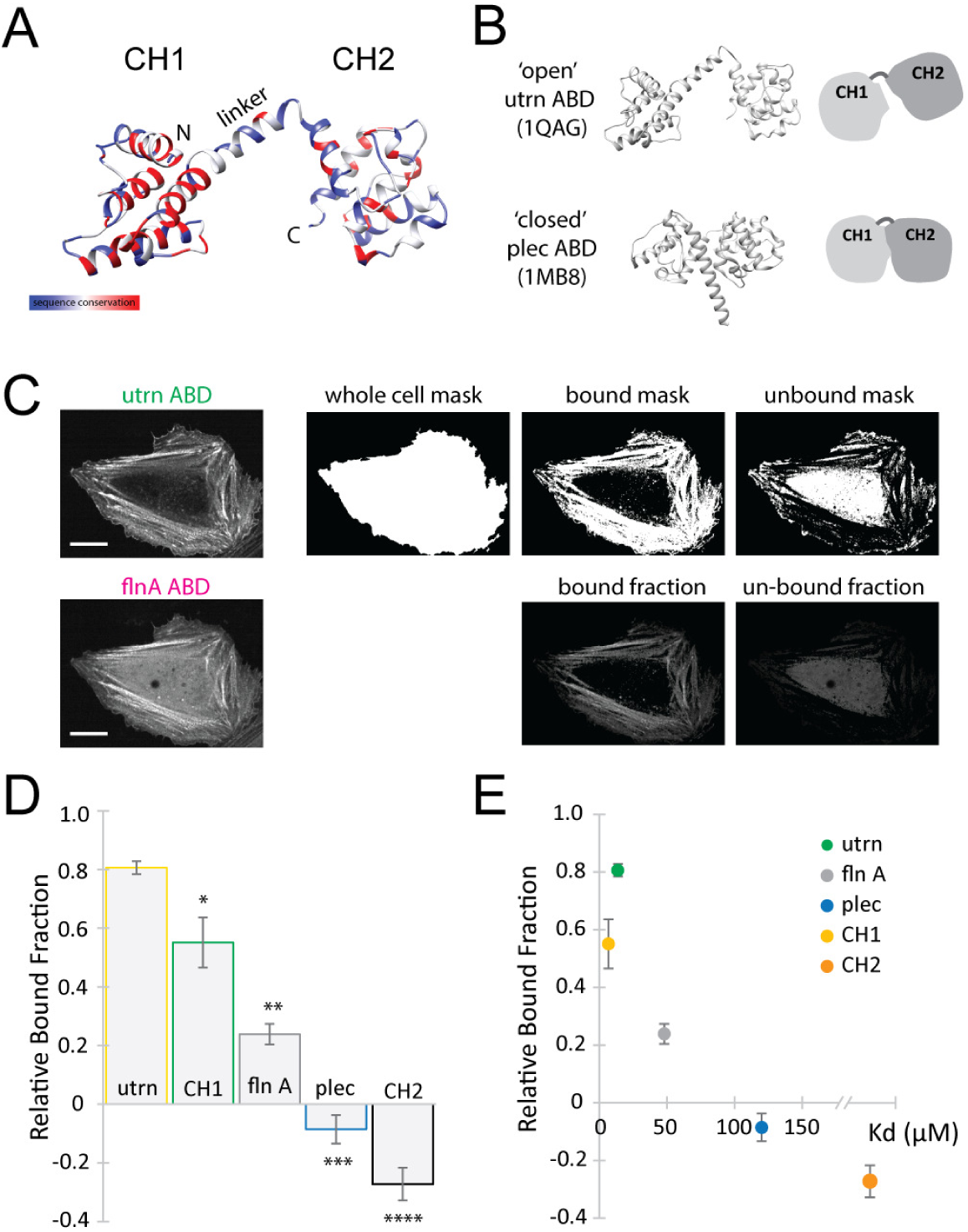
Measurement of bound fraction in live cells correlates with binding affinity *in vitro*. (A) Ribbon diagram of the actin-binding domain of utrophin (1QAG). Colored for sequence similarity between utrophin, filamin and plectin from blue to red. (B) Open and closed conformation of CH1-CH2 from the actin-binding domain of utrophin (1QAG, open) and the actin-binding domain of plectin (1MB8, closed). (C) Method to quantify the relative bound fraction of proteins using live cell imaging. Example images for the actin-binding domain of filamin A. The utrophin ABD channel is used to generate masks for the whole cell, for bound to actin, and for unbound protein, which are then used to calculate intensities in the CH1-CH2 channel of filamin A (bottom row, scale bar 20µm). (D) Relative bound fraction measurements compared to that from utrophin for the actin-binding domains from filamin A (p**<0.05), plectin (p***<0.05), CH1 from utrophin (p*<0.05) and CH2 from utrophin (p****<0.05). (E) Comparison of measurements of relative bound fraction in cells with *in vitro* binding affinity measurements. The K_d_ for CH1 = 6µM (Singh *et al*, 2014), the K_d_ for CH2 > 1000µM (Singh *et al*, 2014), the K_d_ for flnA = 47µM (Ruskamo & Ylänne, 2009), the K_d_ for plectin ≈ 120µM and the K_d_ for utrn = 13.8µM (this study).

The ability for small differences in sequence to have a significant impact on function is particularly clear in disease-associated CH1-CH2 mutations. Mutations to the CH1-CH2 domain of α-actinin-4 are associated with Focal Segmental Glomerulosclerosis, a kidney disorder (Weins *et al*, 2007; Ehrlicher *et al*, 2015; Feng *et al*, 2018), while mutations to the CH1-CH2 domain of filamin’s isoforms and dystrophin are associated with Skeletal Dysplasia (Clark *et al*, 2009; Krakow *et al*, 2004), Muscular Dystrophy (Norwood *et al*, 2000), and the migratory disorder Periventricular Nodular Heterotopia (PVNH) (Parrini *et al*, 2006).

Many of these diseases are a consequence of single point mutations that can result in either loss-of-function (decreased affinity for f-actin) or gain-of-function (increased affinity for f-actin). For example, the K255E mutation in α-actinin-4 (Lee *et al*, 2008) and the M251T mutation in filamin C increase actin-binding affinity (Duff *et al*, 2011), while missense mutations to filamin A in PVNH decrease binding affinity (Iwamoto *et al*, 2018). The K255E mutation to α-actinin-4 is associated with the disruption of an interaction between a tryptophan on the CH1 domain and a cation on CH2. This interaction is highly conserved among CH1-CH2 domains and is proposed to dominate inter-CH domain interactions and affect affinity by latching the domains into a compacted or ‘closed’ configuration (Bañuelos *et al*, 1998; Iwamoto *et al*, 2018; Galkin *et al*, 2010) (Fig 1B). Disruption of this interaction is proposed to allow the domains to adopt an open configuration upon binding to actin, reducing steric clash between CH2 and f-actin and increasing actin-binding affinity (Galkin *et al*, 2010). Physiologically, such increases in actin-binding affinity can cause excessive bundling and crosslinking of the cytoskeleton, compromising cellular function (Weins *et al*, 2007; Avery *et al*, 2017b) and result in changes to the physical properties of the cytoskeleton (Harris *et al*, 2018; Fletcher & Mullins, 2010; Yao *et al*, 2011; Moeendarbary & Harris, 2014). Consequently, precise tuning of CH1-CH2 affinity for f-actin appears to be critical for proper organization and function of the actin cytoskeleton.

Here, we focus on the actin-binding protein utrophin and show that its CH1-CH2 domain affinity for f-actin can be both increased and decreased by perturbations that affect the degree to which it can adopt an ‘open’ or ‘closed’ configuration in solution and reach a bound state through displacement of CH2 upon actin binding. We find that mutations distinct from the well-studied cation-π interaction impacted the affinity of utrophin CH1-CH2, showing that diverse modifications can alter the steric clash between CH2 and f-actin and allow tuning of CH1-CH2 domain affinity. We find that point mutations at the CH1-CH2 interface and replacing the utrophin CH1-CH2 linker domain with an unstructured linker lead to increased affinity, while a PEG modification of the CH2 domain that adds molecular size leads to decreased affinity. These perturbations are consistent with a model in which the degree to which the CH1 and CH2 domains can adopt ‘open’ or ‘closed’ conformations in solution regulates affinity to f-actin. Interestingly, we also find that the N-terminal region of CH1, which was recently shown to affect f-actin binding affinity (Avery *et al*, 2017a; Singh *et al*, 2017; Iwamoto *et al*, 2018), is sufficient to alter the subcellular localization of utrophin’s CH1-CH2 domain in live cells, even compared to mutants with similar bulk affinity. The ability of small sequence changes in CH1-CH2 domains to not only increase and decrease affinity but also alter sub-cellular localization provides new insight into disease-associated mutations and how spatial organization of the actin cytoskeleton is regulated.

## RESULTS

### Measurement of CH1-CH2 domain binding in vitro and in live cells

The overall goal of this study is to understand how changes to a CH1-CH2 actin-binding domain alter its affinity to f-actin, using utrophin as an initial model. To characterize the binding of CH1-CH2 domains to f-actin, we used two complementary approaches: i) traditional *in vitro* co-sedimentation assays using purified proteins and ii) live cell assays in which the relative fraction of protein bound to the actin cytoskeleton is quantified. While co-sedimentation is a standard method for obtaining bulk affinity measurements, a live cell assay offers a more rapid and convenient, albeit less quantitative, way to screen mutants for differences in enrichment on the actin cytoskeleton, as well as for localization to specific structures. We first sought to test whether CH1-CH2 binding assays in live cells would produce results consistent with traditional co-sedimentation assays.

To measure the bound fraction of a fluorescent protein expressed in live cells, we developed a custom image analysis approach based on relative labeling of the actin cytoskeleton. Actin was imaged by expressing the actin-binding domain of utrophin, which is commonly used as a live-cell label of f-actin (Burkel *et al*, 2007), fused to GFP. In a second fluorescence channel, the actin-binding domain-of-interest fused to mCherry was imaged. The average amount of the domain of interest bound to actin was then quantified using the utrophin channel to differentiate between bound and unbound populations (see Materials and Methods). An example showing the actin-binding domain of filamin A compared to utrophin ABD is given in Fig 1C.

We first quantified the binding of CH1 and CH2 domains alone in live cells and compared to previous measurements of CH1 and CH2 affinity. Affinity of tandem CH1-CH2 domains for f-actin is known to primarily arise from the CH1 domain, as CH2 alone cannot bind to actin (Singh *et al*, 2014). We separately expressed the minimal CH1 and CH2 domains from utrophin fused to mCherry (Fig S1). The isolated CH1 domain of utrophin had a high relative bound fraction (0.55±0.09, p*<0.05) (Fig 1D), although the isolated domain appeared partially insoluble, aggregating within cells (Fig S1 A,B), consistent with previous observations about its stability in vitro (Singh *et al*, 2014). The isolated CH2 domain was soluble but distributed throughout the cytosol (Fig S1 C) with a low relative bound fraction (-0.27±0.06, p****<0.05) (Fig 1D), implying that it alone has minimal actin-binding activity. These measurements of relative bound fraction in live cells are consistent with *in vitro* affinity measurements for the isolated CH domains for utrophin (Fig 1E, (Singh *et al*, 2014)) and can be obtained rapidly for screening purposes.

### Relative affinity of CH1-CH2 domains for f-actin can be detected in live cells

We next compared the binding of native CH1-CH2 domains in our live cell assay with co-sedimentation affinity measurements. In tandem configurations, the CH2 has been shown to act as a negative regulator of f-actin-binding through a steric clash with the actin filament upon engagement of the CH1 with f-actin (Galkin *et al*, 2010). In order to bind with high affinity, the CH1-CH2 conformation is thought to adopt an ‘open’ rather than a ‘closed’ conformation, where the steric interaction of the CH2 with f-actin is reduced (Galkin *et al*, 2010). Native tandem CH domains have been shown to crystalize in a range of different conformations, including an ‘open’ state for utrophin actin-binding domain (utrophin’s ABD) (1QAG (Keep *et al*, 1999), Fig 1B) and a ‘closed’ state for plectin’s ABD (1MB8 (García-Alvarez *et al*, 2003), Fig 1B).

We measured the relative bound fraction of the CH1-CH2 domains of utrophin (Fig S1D), filamin A (Fig S1E), and plectin (Fig S1F) in live cells. We found that utrophin had the highest relative bound fraction (0.81±0.02, Fig 1D), while plectin had the lowest (-0.09±0.05, p***<0.05, Fig 1D), characterized by a greater cytoplasmic signal (Fig S1 F). These measurements are consistent with previous data showing that utrophin resides in an ‘open’ conformation in solution, while plectin resides in a ‘closed’ conformation that presents a significant steric barrier to interactions with f-actin (Galkin *et al*, 2010; Lin *et al*, 2011; García-Alvarez *et al*, 2003). To directly compare our relative bound fraction measurements in live cells with bulk affinity measurements, we purified plectin and utrophin ABD’s and measured affinity to f-actin in co-sedimentation assays (Fig 1E, Fig S2). Utrophin’s ABD had a significantly higher binding affinity for actin (K_d_=13.8µM) than that of the plectin construct, which showed little binding over the range of actin concentrations that we tested with our assay (K_d_≈120µM). Together, these results show that a wide range of CH1-CH2 affinities can be captured by measuring relative bound fraction in live cells, though any differences in binding affinity due to actin isoforms could not be assessed.

### Mutations targeting the inter-CH domain cation-π interaction increase the binding affinity of CH1-CH2 domains from plectin but not utrophin

The ‘open’ and ‘closed’ model of tandem calponin homology domain binding to f-actin has focused primarily on the role of a conserved cation-π interaction at the CH1-CH2 interface that latches the CH domains into a ‘closed’ configuration. Typically this interaction is between a highly conserved aromatic residue, e.g. tryptophan, on the CH1 and typically a lysine on the CH2 domain (Borrego-Diaz *et al*, 2006). We wondered whether disrupting this interaction would broadly increase binding affinity, even of CH1-CH2 domains like utrophin’s, which is already considered to be in an ‘open’ configuration.

To test this, we made mutations to the CH2 domains of utrophin (K241E, Fig S1 G), filamin A (E254K, Fig S1 H), and plectin (K278E, Fig S1 I) that are predicted to lie at the interdomain interface and measured the resulting bound fractions in live cells (Fig 2A). Consistent with the ‘open’ and ‘closed’ model of CH1-CH2 binding, these mutations increased the binding of the filamin A (0.64±0.05, p<0.05) and plectin domains (0.62±0.03, p**<0.05) to f-actin, resulting in a bound fraction closer to that of the native utrophin CH1-CH2. However, the mutation of the equivalent residue on utrophin ABD had no effect on the apparent bound fraction, consistent with the idea that this domain already exists in an ‘open’ conformation relative to filamin and plectin ABDs (Fig 1A,B 0.84±0.02, p=0.08) (Lin *et al*, 2011). We made co-sedimentation measurements of the plectin K278E mutant that confirmed its bulk binding affinity increased significantly (K_d_=45.1µM).

**Figure 2:**
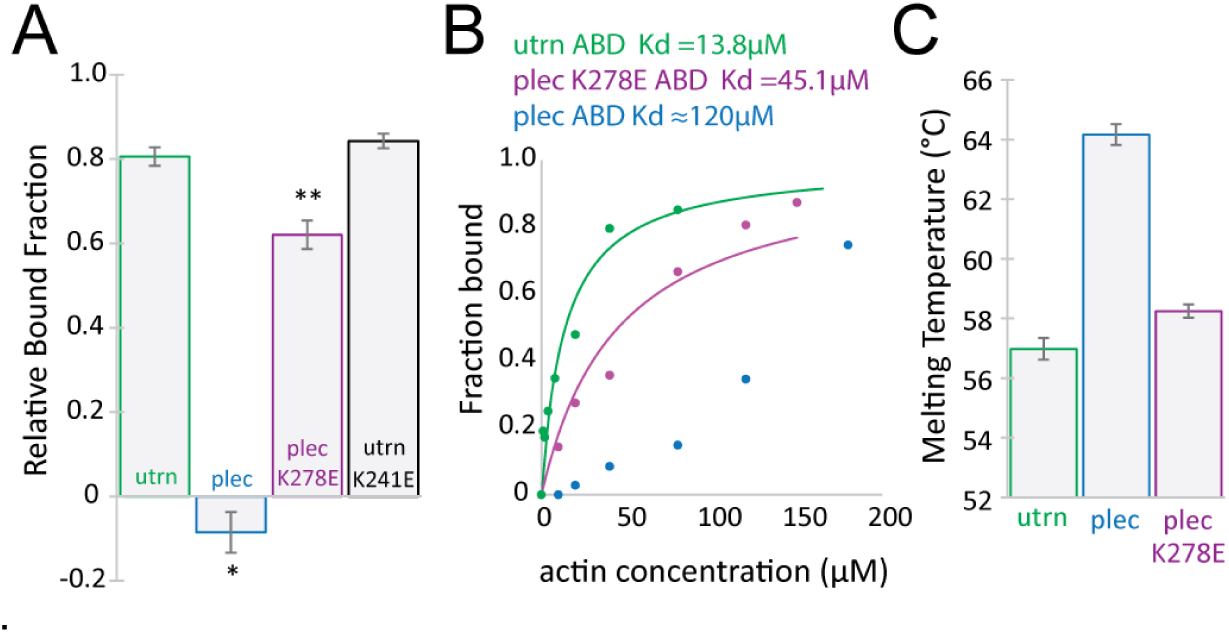
Binding affinity to f-actin depends on conformation and inter-CH domain interactions. (A) Relative bound fraction measurements for the mutants of plectin (K278E, p**<0.05) and utrophin (K241E, p=0.08). (B) Binding curves of actin-binding domains to f-actin. Low binding is observed for plectin over the range of concentrations tested, implying a low affinity interaction. The mutation K278E restores binding affinity. (C) Melting temperatures for the actin-binding domain of plectin and K278E mutant of plectin’s actin-binding domain.

To test inter-domain interactions in a different way, we measured melting temperature as a proxy for domain stability (Avery *et al*, 2017a; Singh *et al*, 2014; Singh & Mallela, 2012). As expected, the melting temperature of the plectin CH1-CH2 (Tm=64.2±0.4°C) was higher than that of the utrophin CH1-CH2 (Tm=57.0±0.4°C, Fig 2C), implying a more compact and stable conformation, while the plectin K278E mutation had a reduced melting temperature in comparison to the native domain (Tm=58.2±0.2°C, Fig 2C), consistent with reduced inter-CH domains.

### Utrophin CH1-CH2 affinity is increased by alternate interface mutations that do not change solution ‘openness’

The mutation that targets the π-cation interaction reduced inter-domain interactions for plectin’s CH1-CH2 domain and increased its affinity for f-actin but had no effect on utrophin’s CH1-CH2 domain. However, disease-associated mutations have been shown to change the affinity of tandem calponin homology domains by several orders of magnitude (Avery *et al*, 2017b), suggesting that other CH1-CH2 interactions could impact binding affinity. To test this, we focused on utrophin’s CH1-CH2 and investigated the contribution of different parts of the ABD to actin binding affinity.

We introduced the point mutations Q33A and T36A to the CH1-CH2 domain of utrophin, locations that are predicted to lie at the CH1-CH2 interface and evaluated f-actin binding. In our live cell assay, the relative bound fraction of this construct remained high (Fig 3A, 0.87±0.05, p=0.23), indicating a high actin-binding affinity. We then measured the mutant’s binding affinity in a co-sedimentation assay and observed a significantly higher actin-binding affinity (K_d_=0.4µM, Fig 3B) than the native utrophin CH1-CH2. Consistent with this increase in affinity, we measured a lower melting temperature (Tm = 54.9±0.3°C, Fig S2D) for the mutant CH1-CH2 compared to the native domain, indicating reduced structural stability. We speculated that destabilization of CH1-CH2 interactions could increase binding affinity by two mechanisms. First, the mutations could shift the solution state of the domain to be ‘more open’ – further reducing steric interactions upon initial binding to f-actin. Secondly, they could make transitioning to the bound state more favorable – in the absence of changes in solution openness.

**Figure 3:**
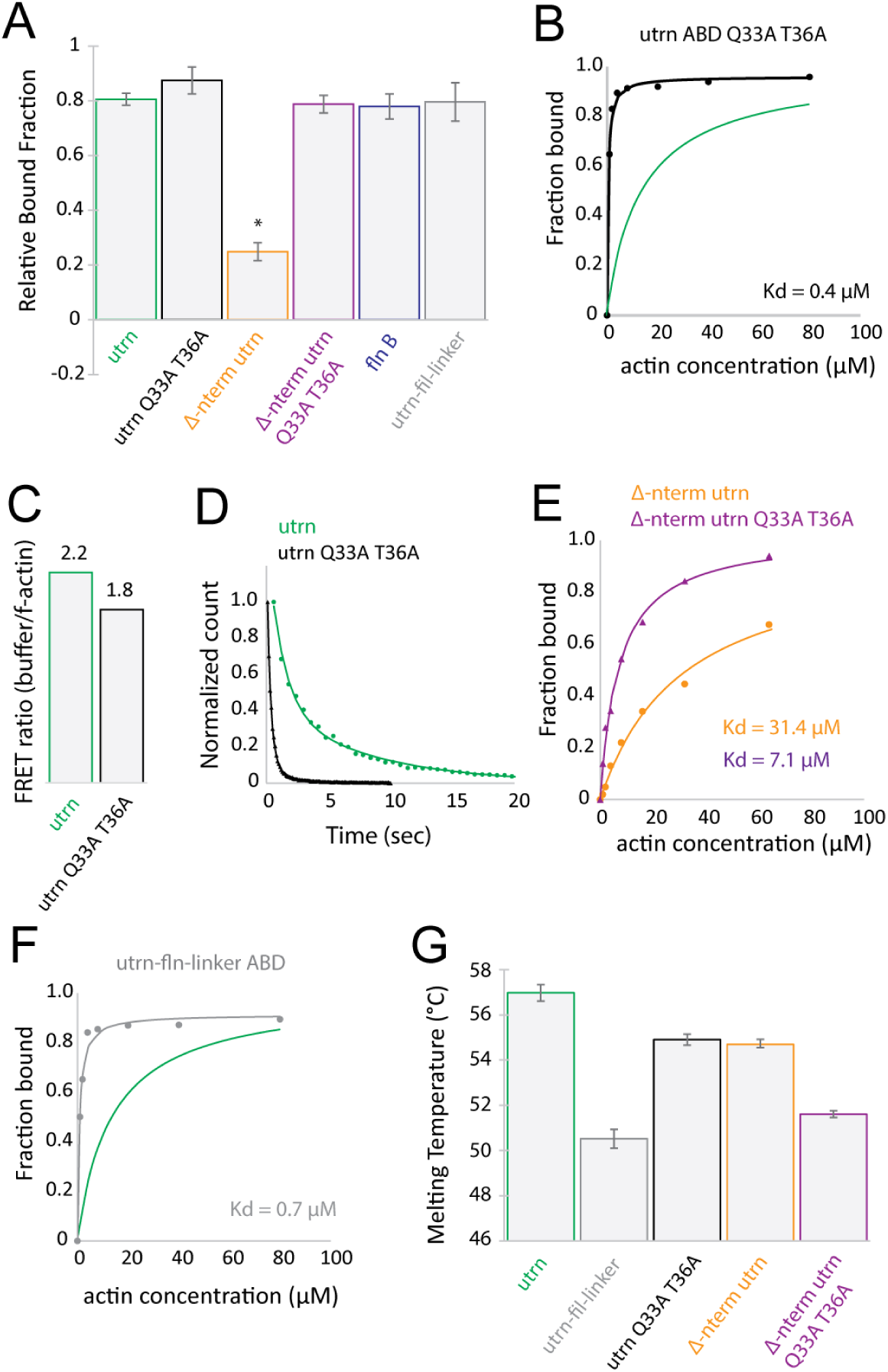
Mutations to the interdomain interface or interdomain linker region result in an increase in binding affinity. (A) Relative bound fraction measurements for interdomain interface mutants (p=0.24), n-terminal deletion (p<0.05), n-terminal deletion with Q33A T36A (p=0.54), the actin binding domain from filamin B (p=0.62) and a chimera of the interdomain linker from filamin A and utrophin’s CH domains (p=0.89). (B) Binding curves for the Q33A T36A mutation (black, K_d_ = 0.4µM). The binding fit for WT utrn is shown in green for comparison. (C) FRET ratio of domains in buffer and in the presence of actin (WT utrn ABD shown in green and Q33A T36A shown in black). (D) Single molecule binding dwell time histograms. (E) Binding curves for Δ-nterm (orange, K_d_ = 31.4µM) and Δ-nterm Q33A T36A (magenta, K_d_ = 7.1µM). (F) Binding curves for the utrn-fln-linker chimera (grey, K_d_ = 0.7µM). The binding fit for WT utrn is shown in green for comparison. (G) Melting temperature measurements for the interdomain linker mutant and the interdomain interface mutant of utrophin’s actin-binding domain.

To investigate whether the interface mutations altered physical properties of CH1-CH2 domains, we measured radius of gyration (Rg), which captures CH1-CH2 ‘openness’, using small angle X-ray scattering (SAXS) (Hura *et al*, 2009) (Fig S3, see Materials and Methods). The plectin ABD construct had the lowest Rg (22.0Å), suggesting a ‘closed’ conformation in solution, while the Rg of utrophin ABD was larger (24.0Å). The utrophin Q33A T36A mutant had a similar Rg to that of WT utrophin (23.5Å), suggesting that the solution ‘openness’ of the domains were similar. However, our SAXS data also indicated a slight increase in flexibility of the domain.

### FRET and single molecule measurements reveal changes in utrophin CH1-CH2 mutant binding kinetics

WT utrophin has been shown to undergo a conformational change when binding to f-actin through an induced fit mechanism (Lin *et al*, 2011). We sought to test whether there was a similar structural change of the Q33A Q36A mutant upon binding to f-actin. We compared opening of the domains using Förster Resonance Energy Transfer (FRET) in the presence and absence of f-actin. To do this, we installed an N-terminal GFP as the donor and engineered a single cysteine at position 168 on the CH2, which we chemically labelled with Alexa 555 maleimide as an acceptor fluorophore. Interestingly, both the WT and mutant domains showed a decrease in FRET in the presence of f-actin (Fig 3C), suggesting that both undergo an induced-fit upon binding to actin.

If the mutant utrophin ABD has similar solution ‘openness’ to WT utrophin ABD based on SAXS measurements and similar reduction in FRET upon f-actin binding, what could give rise to the difference in bulk affinity? We wondered if comparing binding kinetics of the two ABDs could provide further insight. We measured the dwell times of single molecule binding events of WT utrophin and the Q33A T36A mutant with TIRF microscopy (Fig 3D, Fig S4). Interestingly, the mean dwell time of binding events from the Q33A T36A mutant was ~10-fold longer than that of the WT binding domain (T_Q33AT36A_ = 8.49 ± 0.14 sec, T_WT_ = 0.96 ± 0.01 sec). However, the difference in binding affinity measured using co-sedimentation was ~30 fold, suggesting a difference in on-rate of ~3-fold. In summary, the Q33A T36A mutations affected both the on-rate and off-rate of binding.

### Loss of utrophin CH1-CH2 affinity due to N-terminal truncation can be compensated by the incorporation of CH1-CH2 interface mutations

The N-terminal flanking region varies significantly between CH1-CH2 domain proteins, both in sequence and in length (Iwamoto *et al*, 2018; Singh *et al*, 2017). This region has recently been shown to be important for actin-binding affinity, as its deletion in either utrophin (Singh *et al*, 2017) or ß-spectrin (Avery *et al*, 2017a) reduces actin binding. Interestingly, the N-terminal flanking regions from filamin B is significantly shorter than that from ß-spectrin, despite the relatively high reported binding affinity (K_d_~7µM) of filamin B’s ABD for f-actin (Sawyer *et al*, 2009).

To confirm the importance of the N-terminal flanking region in CH1-CH2 affinity for f-actin, we truncated residues 1-27 of the utrophin ABD and expressed the remaining CH1-CH2 domain in live cells (Fig 3A, Fig S5A). This construct (Δ-n-term) had a low bound fraction (0.25±0.03, p<0.05), indicating a reduced binding affinity to F-actin compared to the native utrophin CH1-CH2. This is consistent with previous results reporting the importance of this region for actin-binding affinity (Iwamoto *et al*, 2018; Avery *et al*, 2017a; Singh *et al*, 2017).

We wondered whether it would be possible to compensate for the loss in binding affinity by modifying the inter-CH domain interface of the Δ-n-term construct, as demonstrated above. To test this idea, we introduced the mutations Q33A T36A, which increased the affinity of the native utrophin CH1-CH2 domain into the n-terminal truncation construct (Δ-n-term Q33A T36A) (Fig 3A). Remarkably, this mutation restored the bound fraction of the mutant Δ-n-term to f-actin in live cells (0.79±0.03, p=0.54, Fig 3A, Fig S5B). Consistent with this, the incorporation of mutations to the inter-CH domain interface recovered the binding affinity of the n-terminal truncation as measured by co-sedimentation (Fig 3E, K_d_ Δ-n-term = 31µM, K_d_ Δ-n-term Q33A T36A = 7µM). These findings suggest that some mechanisms controlling CH1-CH2 affinity, including contributions from inter-CH domain interactions and the N-terminal region, contribute to affinity in a separate and additive manner.

### Utrophin CH1-CH2 interdomain linker structure affects binding affinity

Similar to the N-terminal flanking region, the interdomain linker region has a high level of sequence and structural diversity among native CH1-CH2 domain-containing proteins. The linker can be unstructured, as in the case of filamin and plectin (not resolved in the crystal structures of filamin A 2WFN (Ruskamo & Ylänne, 2009) or plectin 1MB8 (García-Alvarez *et al*, 2003)), or it can be helical, as in the case of utrophin (1QAG (Keep *et al*, 1999)) and dystrophin (1DXX (Norwood *et al*, 2000)). We postulated that the interdomain linker region could have a role in regulating CH1-CH2 domain ‘openness’ in solution and thereby its affinity to f-actin. To test this, we generated chimeras containing the CH1 and CH2 domains from utrophin but the linker region from filamin A. In our live cell assays, this chimeric protein had a high relative bound fraction (0.80±0.07, p=0.89) (Fig 3A), indicating a high affinity for f-actin. We next expressed and purified this construct and found that the actin-binding affinity based on co-sedimentation was significantly higher (K_d_=0.7µM) (Fig 3F) than that of the WT utrophin CH1-CH2. We also found that the chimeric protein had a significantly lower melting temperature (Tm = 50.5±0.4°C) (Fig 3G), indicating that the filamin A unstructured linker caused a ‘more open’ configuration of the tandem CH domain in solution than the WT utrophin CH1-CH2.

To further test the effect of the linker on properties of the CH1-CH2, we measured the radius of gyration (Rg) and flexibility of the linker chimera CH1-CH2 using small angle X-ray scattering (SAXS) (Hura *et al*, 2009) (Fig S3, materials and methods). Compared to the ‘closed’ plectin CH1-CH2 domain (Rg = 22Å) and ‘open’ utrophin CH1-CH2 domain (Rg = 24Å), the utrophin-filamin-linker chimera had the largest Rg of the constructs we tested (41Å), indicating that the unstructured linker from filamin A allowed the domain to adopt a conformation with a large separation between the CH1 and CH2 in solution. This construct was also the most flexible, potentially reducing any steric clash between the CH2 and actin filament upon binding, resulting in a higher affinity toward f-actin.

### Increasing the steric interaction of utrophin CH2 domain with f-actin can decrease f-actin binding affinity

As seen in our measurements above and previous studies (Galkin *et al*, 2010), ‘opening’ of the CH1-CH2 interface is believed to reduce steric clash with the actin filament and provide increased accessibility for CH1 to the f-actin surface. We wondered if it would be possible to reduce the f-actin affinity of CH1-CH2 domains that are in an ‘open’ configuration in solution (e.g. native utrophin CH1-CH2, disease associated gain-of-function mutants, utrophin interface mutants, or utrophin-filamin-linker chimera) by modifying the CH2 domain to increase steric clash. Since CH2 alone has little to no binding interaction with f-actin (Fig 1D), we postulated that its steric interaction with actin could be increased by adding biologically-inert bulk that simply increases the molecular size of the domain.

We increased the size of the CH2 domain of the WT utrophin CH1-CH2 domain by conjugating a small PEG molecule to the surface of the domain (Fig 4A). Specifically, we mutated a serine, S158, to cysteine and performed a conjugation reaction with maleimide-750Da PEG, which has an approximate Rg of 1nm. We found that the unconjugated mutant had a similar binding affinity to WT utrophin CH1-CH2 (K_d_=12.5µM, Fig 4B), but the PEG-conjugated utrophin CH1-CH2 had a reduced binding affinity (K_d_=53.8µM, Fig 4B). This result highlights that simply increasing the physical size of CH2 can the reduce gains in affinity that arise from ‘open’ conformation domains, supporting the idea that the CH2 steric clash with f-actin indeed modulates CH1-CH2 affinity.

**Figure 4:**
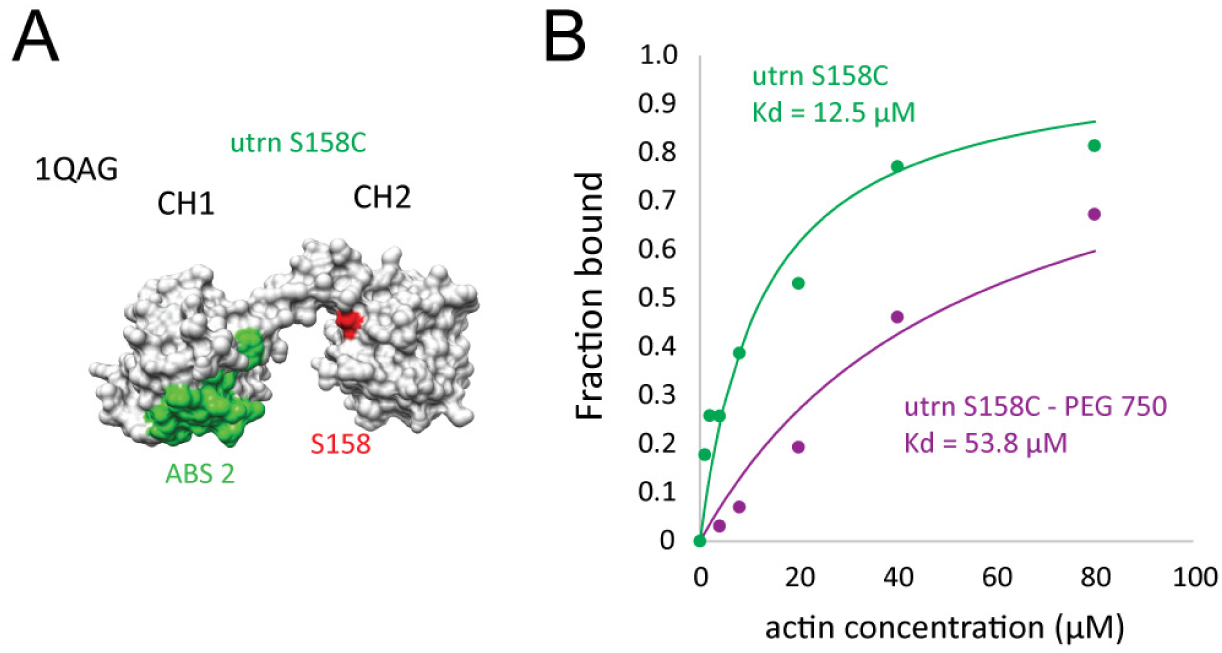
Binding affinity of CH1-CH2 to f-actin can be reduced by increasing CH2 size. (A) Surface model of the actin-binding domain of utrophin (1QAG). The f-actin-binding surface on CH1, ABS2 (Iwamoto *et al*, 2018), is shown in green, and residue S158 on CH2 is shown in red, which was mutated to cysteine and used for PEG conjugation. (B) Binding curves for the utrophin S158C mutant (green) and the PEG750 conjugated mutant (magenta). This size increase caused a change in binding affinity from K_d_=12.5µM to K_d_=53.8µM.

### CH1-CH2 domain subcellular localization is affected by the N-terminal region, independent of affinity

Our live cell assay for CH1-CH2 binding allows us to screen not only for binding affinity but also sub-cellular localization to different actin structures. We quantified differences in localization by calculating the correlation coefficient relative to native utrophin CH1-CH2 and by measuring the bound amounts of proteins on different actin structures (e.g. stress fibers vs peripheral actin networks). We first examined the utrophin-filamin CH1-CH2 linker chimera and found that its sub-cellular localization differed significantly from the WT utrophin CH1-CH2 domain. Specifically, it had a low correlation coefficient over the whole actin cytoskeleton (0.54±0.04, p<0.05, Fig 5A) and was comparatively depleted from the cell periphery (Fig S5). However, as reported above, the CH1-CH2 linker chimera affinity was significantly larger than that of the WT utrophin ABD. This large difference in affinity makes it difficult to conclusively decouple the contribution of affinity on subcellular localization to different actin structures.

**Figure 5:**
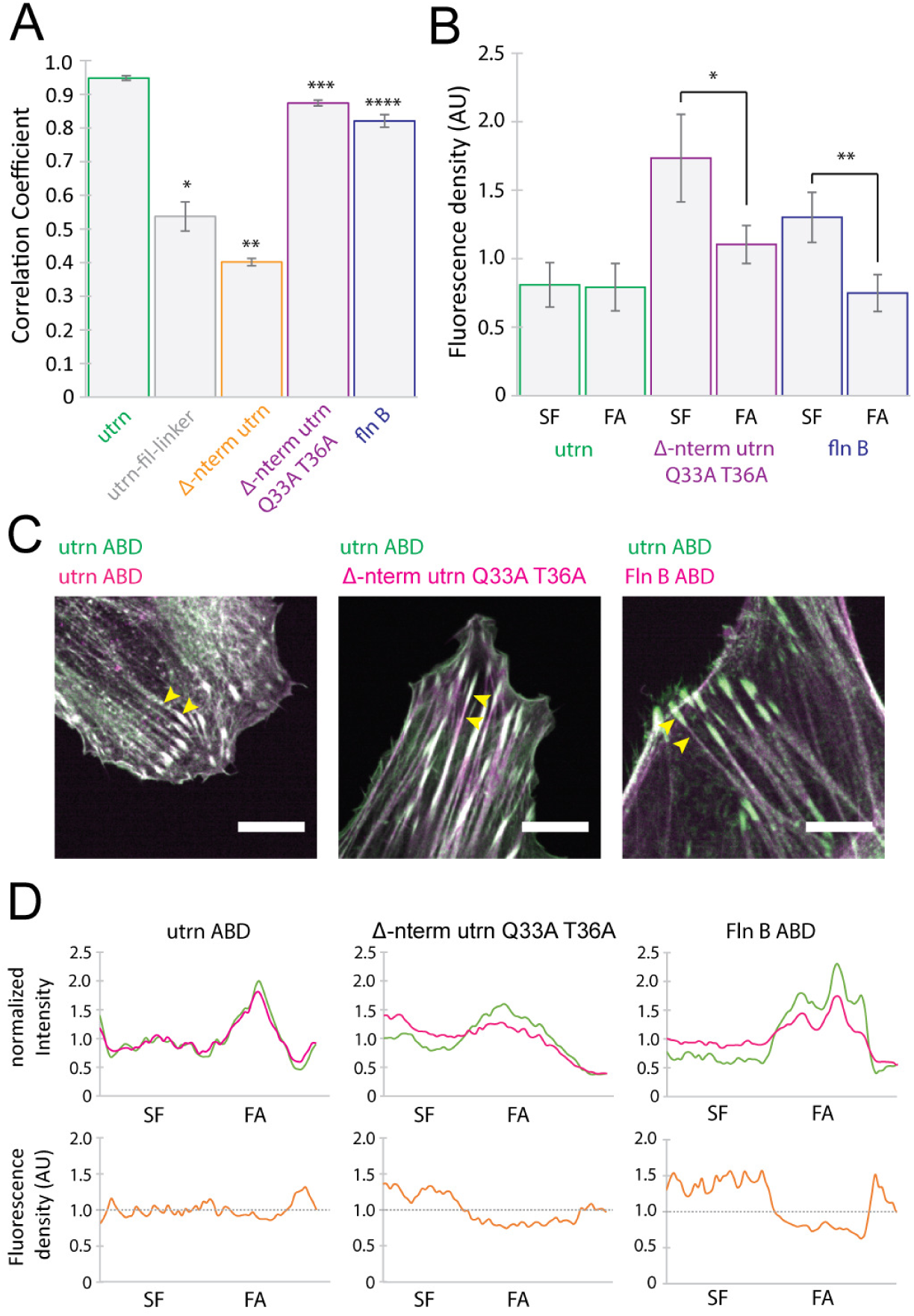
The N-terminal flanking region plays a role in CH1-CH2 localization to different actin structures. (A) Measurements of whole cell correlation coefficients for utrophin (utrn), utrophin-filamin-linker (p*<0.05), Δ-nterm (p**<0.05), Δ-nterm Q33A T36A (p***<0.0.5) and filamin B (p****<0.05) relative to WT utrophin ABD. (B) Measurement of protein-binding density on different actin structures (stress fibers and focal adhesions) show significant differences in binding localization for both the Δ-nterm Q33A T36A construct (magenta, p*<0.05) and the CH1-CH2 from filamin B (blue, p**<0.05). (C) Representative images of the different constructs. (D) Line-scans sectioning a stress fiber terminating in a focal adhesion showing the intensity in each channel and the density of the construct of interest (magenta) across the line-scan. Scale bars are 10µm.

To compare subcellular localization of different CH1-CH2 domains with similar affinity, we turned to the utrophin ABD N-terminal truncation with interface mutations that we introduced previously (Δ-n-term Q33A T36A). Interestingly, the subcellular localization of the mutant was significantly different from WT utrn CH1-CH2 domain (Fig 5B, Fig S6). The Δ-n-term Q33A T36A mutant displayed a moderate correlation with the WT utrophin ABD over the whole actin cytoskeleton (0.87±0.03). Subcellularly, this mutant was distributed evenly on stress fibers and focal adhesions, while the WT utrophin CH1-CH2 was comparatively more enriched in focal adhesions (Fig 5C,D).

Finally, we investigated the subcellular localization of filamin B’s CH1-CH2 domain, which has a short N-terminal region and a K_d_ = 7µM, which is on the same order as WT utrophin CH1-CH2 (K_d_=13.8µM). Surprisingly, this domain showed preferential localization to stress fibers and was comparatively reduced at focal adhesions (Fig 5C,D), similar to that of the utrophin construct with N-terminal truncation (Δ-n-term Q33A T36A). These differences in localization were not a result of differences in the dynamics of the proteins as measured by Fluorescence Recovery After Photobleaching (FRAP) (Fig S7A), indicating that the N-terminal region may play a key role in modulating specificity of CH1-CH2 domains for different actin structures, independent of bulk differences in actin-binding affinity and dynamics.

## DISCUSSION

Diseases involving actin-binding proteins with CH1-CH2 mutations that exhibit a gain-of-function (increased actin-binding affinity) are associated with increased ‘openness’ of the domains. This often involves the disruption of a conserved cation-π interaction that is proposed to dominate inter-CH domain interactions and hold the two globular CH domains in a compact configuration.

Consistent with this, we found that disruption of the conserved cation-π interaction increased the binding affinity of plectin to be more like that of utrophin, which is thought to reside in an ‘open’ configuration. By making chimeras of utrophin’s CH1 and CH2 with the linker region from filamin A, we have shown that the interdomain linker region can impact binding affinity by altering the ‘openness’ of CH1 and CH2. Previous work has compared chimeras prepared from the CH domains of utrophin and the interdomain linker region of dystrophin, which did not significantly change binding affinity (Bandi *et al*, 2015). Importantly, however, the linkers from utrophin and dystrophin both have a helical structure, which might not be expected to alter domain ‘openness’. By introducing the unstructured linker from filamin A, we observed a large increase in binding affinity. In this configuration, the CH2 can presumably move away more freely from f-actin, thereby reducing any possible steric interactions with the filament that would hinder CH1 binding. This increased ‘openness’ is consistent with the large Rg of the chimera in solution observed in our SAXS measurements. Furthermore, mutations to the linker region of utrophin’s native CH1-CH2 that are predicted to destabilize its helical structure had a similar behavior to the filamin linker-utrophin chimera when expressed in cells (Fig S5C).

If steric clash between CH2 and f-actin is reduced when CH1-CH2 domains are in ‘open’ configurations in solution, then increasing steric clash should reduce binding affinity. We directly test this idea by adding size to the CH2 domain and measuring binding to f-actin. After conjugating a biochemically inert PEG molecule (Rg~1nm) to the CH2 domain, we find that overall affinity of the domain is reduced ~5 fold. Interestingly, this concept could present a potential therapeutic approach for diseases that result in gain-of-actin-binding-function, where a molecule of a specific size would target the CH2 domain in order to increase the steric interaction between CH2 and f-actin, thereby reducing binding affinity. The novelty of this approach is that the interaction between CH1 and f-actin itself does not need to be disrupted, meaning that overall affinity can be reduced without completely abolishing it by blocking or antagonizing the CH1 to actin binding interface.

Interestingly, we found that additional mutations to the CH1-CH2 interface of utrophin distinct from the well-studied cation-π interaction, caused an increase in actin-binding affinity without altering its solution ‘openness’. In our measurements, both WT utrn ABD and the Q33A 36A mutant had similar Rg values and both underwent a conformational change when binding to actin, as measured by FRET. These observations suggest that mutations to the inter-CH domain interface make it easier for the protein to undergo a conformation change when binding, characterized by a ~3-fold increase in on-rate, while not having a dramatic effect on their ‘openness’ in solution. Furthermore, we observed a change in binding off-rate of ~10 fold for the Q33A T36A mutant. We speculate that the large difference in off-rate implies that reduced inter-CH domain interactions allow the domain to adopt a high-affinity state when bound to f-actin, potentially through reduced steric interactions between CH2 and f-actin. This notion is consistent with measurements that have shown that the CH domains from dystrophin and utrophin can adopt a range of conformations in solution, only some of which are potentially compatible with f-actin binding (Fealey *et al*, 2018).

Finally, we observed that the N-terminal flanking region prior to CH1 appears to affect CH1-CH2 domain localization to specific actin structures, independent of binding affinity. By truncating the N-terminal flanking region of utrophin (which reduces affinity) and introducing CH1-CH2 interface mutations (which increases affinity), we were able to create a construct with similar binding affinity to WT utrophin CH1-CH2 (Fig 3E, Fig 5A) but with significantly different subcellular localization (Fig 5 B-D). The change in localization is not the result of kinetic differences in binding, which have been proposed for the localization of myosin to the rear of migratory cells (Maiuri *et al*, 2015), as the kinetics of both ABDs were similar when measured by FRAP (Fig S7 A).

These results indicate that CH1-CH2 domains could influence both the binding and the localization of full-length proteins that contain them. For example, the isoforms filamin A and filamin B have high sequence identity both across both the full-length protein (~68%) and within the CH1-CH2 domain (~75%), but the minimal actin binding domains have different affinities for f-actin and different localizations, as well as different cellular functions. Filamin A plays a critical role in maintaining cortical mechanical integrity, but the presence of filamin B is not sufficient to compensate for the absence of filamin A in blebbing melanoma cells (Biro *et al*, 2013). Filamins also function as signaling scaffolds, interacting with more than 30 different proteins. Genetic mutations to each isoform are linked with specific filaminopathies suggesting distinct protein interactions between isoforms (Feng & Walsh, 2004). Many genetic mutations that result in filaminopathies are clustered within the actin-binding domain, and it is interesting to speculate that changes in localization could also result in differences in intracellular signaling. One region of increased diversity between these proteins is in the N-terminal flanking region. When we express the actin-binding domains from different filamin isoforms in live cells we observe different binding characteristics. A chimera of the filamin A N-terminal flanking region with CH1 and CH2 domains from filamin B partly increased its subcellular localization to focal adhesions, but some differences in localization (compared to WT utrophin ABD) could still be observed (Fig S8). This result implies that the N-terminal flanking region is indeed important for affinity but also for subcellular localization of the domain. The combination of inter-CH domain and N-terminal interactions therefore create a versatile range of actin-binding properties, including affinity to f-actin and localization to specific actin structures. While CH1-CH2 domains from different proteins share similarities in structure and sequence, small differences can be significant, affecting both binding affinity and localization and highlighting why disease-associated point mutations can have such a detrimental impact.

## MATERIALS AND METHODS

### Cell Culture

HeLa and HEK293T cells were cultured at 37°C in an atmosphere of 5% CO_2_ in air in DMEM (Life Tech, #10566024) supplemented with 10% FBS (Life Tech, #16140071) and 1% Penicillin-Streptomycin (Life Tech, #15140122). HEK293T cells were passaged using 0.05% trypsin and HeLa cells with 0.25% trypsin.

### Generation of constructs and cell lines

To visualize the localization of different constructs with respect to utrn ABD, single expression and bi-cistronic expression plasmids were generated for transient transfection, and two separate virus plasmids were generated for creating stable cell lines. PCS2+ GFP-UtrCH was a gift from William Bement (Addgene plasmid # 26737) (Burkel *et al*, 2007). cDNA for generating constructs to image WT CH domains were either amplified using PCR or synthesized directly (Integrated DNA Technologies) and inserted into the desired vector using Gibson assembly. The actin-binding domain of human filamin A corresponds to residues 1-278, and we used a similar construct to that of Garcia Alvarez et al. for the actin-binding domain of plectin a.a. 60-293 (Garcí́a-Alvarez *et al*, 2003). For dual expression, a cleavable peptide was introduced to the c terminus of GFP-UtrCH followed by mCherry fused to the actin-binding domain of interest (Kim *et al*, 2011). Transient transfections were performed using Effectene (Qiagen, #301425), following the manufacturers stated protocol and imaged 24 hours after transfection. For generating stable cell lines, GFP-UtrCH and the construct of interest fused to mCherry were cloned into Lentiviral plasmid pHR. Lentiviruses were then generated through 2^nd^ generation helper plasmids and their transfection into HEK293 cells for packaging. Lentiviral supernatants were collected 48-72 hours after transfection, filtered using a 0.4um filter, and used directly to infect the target cell line in a 1:1 ratio with normal culture media.

### Protein purification and labelling

Actin was purified from rabbit muscle acetone powder (Pel Freez Biologicals, #41995-1) according to Spudich & Watt, 1971. Actin was stored in monomeric form in G-buffer (2mM Tris-Cl pH 8.0, 0.2 mM ATP, 0.5 mM TCEP, 0.1 mM CaCl_2_) at 4°C. Petm60-Utr261 was a gift from Peter Beiling (Bieling *et al*, 2017). Utrn ABD and its associated mutants were expressed recombinantly in E. coli BL21 (DE3) pLysS (Promega, #L1191) and purified using affinity chromatography followed by gel filtration. Proteins were stored in 20 mM Tris-Cl pH 7.5, 150 mM KCL, 0.5 mM TCEP (GF-buffer) and 0.1 mM EDTA in the presence of 20% glycerol. Utrophin ABD and plectin ABD sequences included a KCK linker (GGSGKCKSA) on the C terminus for labelling. Proteins were labelled using either Alexa 555 and Alexa 488 maleimide dye (Life Technologies, #A22287 & #A20346) at the cysteine site in the KCK linker region. Briefly, proteins were reduced in 5mM TCEP for 30 minutes and then buffer exchanged over a desalting column into GF-buffer without TCEP. Labelling was performed at 4°C with a ~5-fold molar excess of dye overnight. The reaction was then quenched with DTT and the excess dye removed by gel filtration. A typical labelling ratio was ~75%. The actin-binding domain of plectin was purified and labelled using the same method as for utrophin, but with a reduced labelling time to yield a similar labelling ratio to utrophin. For PEG-conjugated utrophin constructs, a similar purification and labelling strategy was used but with EGFP fused to the n-terminus of the domain so that single cysteine mutants could be used for labelling. 750 Da PEG-maleimide (Rapp Polymere) was conjugated to cysteine residues on utrophin using the same labelling strategy described above.

### Spinning Disc Confocal Imaging

Fluorescent proteins were imaged using the following excitation and emission: GFP was excited at 488 nm and emission was collected at 525 nm, mCherry was excited at 543 nm, and emission was collected at 617 nm. Live imaging experiments were performed in normal cell culture media using an OKO labs microscope stage enclosure at 37°C in an atmosphere of 5% CO_2_. Cells were imaged on glass bottomed 8 well chambers that had been coated with 10µg/ml fibronectin. Dual color images were used to measure differences in protein binding and localizations (Belin *et al*, 2014).

### Relative bound fraction measurements and image difference mapping

A custom-written MatLab routine was used to calculate the relative bound fractions of different actin-binding domains, the difference maps and correlation coefficients. Briefly, images of cells expressing GFP-utrn were thresholded and binarized to generate masks of the whole cell and actin cytoskeleton. Holes within the binary image mask were filled and this was used as an outline of the cell footprint. To generate a mask for the unbound fraction the complementary image of the actin mask was taken, which included pixels only within the cell footprint (Fig 1C). Average pixel intensity measurements (Ī) were then made using the two masks and the relative bound amount calculated from the following equation 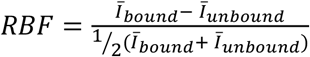. In some instances of very low binding, *RBF* was less than zero. This arises due to the geometry of the cell where higher intensities are gathered from the cell body where there is more cytoplasmic signal, in comparison to actin rich regions that are often thin and flat at the cell periphery. *RBF* is a convenient and relative measure when averaging over many cells to rule out large contributions from cell geometry or f-actin abundance. For comparing the localization of different actin-binding domains, the same masking method was used to make measurements of intensity of the CH1-CH2 of interest and utrn ABD. The images were normalized to their maximum value and the utrn ABD image values subtracted from the CH1-CH2 image to give the difference map (Fig 5B). Pearson’s correlation coefficient was calculated from the image values to quantify whole cell differences in localization. To measure the relative amounts of protein bound to different actin structures (stress fibers, and ocal adhesions), local pixel value measurements were made, background subtracted and normalized to the f-actin abundance within that structure (using the utrn ABD channel as a reference for f-actin).

### Fluorescence Recovery After Photobleaching measurements

To measure the recovery rate of different proteins in live cells we performed fluorescence recovery after photobleaching experiments (FRAP). Measurements were made using a Zeiss LSM 880 NLO Axio Examiner using a 20x dipping objective. Cells were plated onto 6cm plastic bottomed dishes (falcon) 24 hours prior to experiment. Cells were imaged for one frame; a small circular region ~1µm in diameter was then bleached and imaged with a frame rate of 0.95 sec/frame to monitor the fluorescence recovery. Images were analyzed using the approach of Phair et al. (Phair *et al*, 2003). To calculate the proteins’ dynamics, the initial rate of recovery was measured, which is independent of the bleaching fraction or immobile fraction, in contrast to the fluorescence recovery half time.

### Circular Dichroism (CD) and melting temperature measurements

CD wavelength scans (250 to 200 nm) and temperature melts (25°C to 80°C) were measured using an AVIV model 410 CD spectrometer. Temperature melts monitored absorption signal at 222 nm and were carried out at a heating rate of 4°C/min. Protein samples were prepared at ~7.5 µM in PBS in a 0.1 cm cuvette. Melting temperature data was fitted using the equation 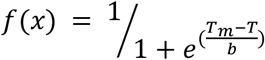 where *T*_*m*_ is the melting temperature and *b* is the slope parameter and *T* is the temperature.

### Small Angle X-ray Scattering measurements

Proteins were exchanged into SAXS buffer (20 mM Tris-Cl pH 7.5, 150 mM KCl, 0.5 mM TCEP and 0.1 mM EDTA) using a 10kDa MWCO Zeba spin desalting column (Thermo Scientific). Corresponding blanks were prepared by diluting flow-through from spin columns into appropriate buffers at the same dilution. Samples were prepared at concentrations of 3-8 mg/mL. SAXS measurements were made at SIBYLS 12.3.1 beamline at the Advanced Light Source. The light path is generated by a super-bend magnet to provide a 1012 photons/sec flux (1 Å wavelength) and detected on a Pilatus 3 2M pixel array detector. Data from each sample was collected multiple times with the same exposure length, generally every 0.3 seconds for a total of 10 seconds resulting in 30 frames per sample. Data was analyzed using the Scatter software.

### Actin Filament Binding Assay

Filamentous actin was prepared by polymerizing β-actin at 162 µM for 1.5 hr at room temperature. Various concentrations of F-actin were then combined with a constant concentration of fluorophore-labeled actin-binding domain (either 100 nM for utrn ABD, utrn-fil-linker ABD, utrn Q33A T36A ABD or 1 µM plectin ABD, and plectin K278E ABD) in Buffer F. Sub stoichiometric concentrations of actin-binding domains were used in all experiments, such that the assumption of [F-actin]_total_ ≈ [F-actin]_free_ was valid. After incubation at room temperature for 30 min, F-actin and bound actin-binding domain were pelleted at 150,000 x *g* for 60 min at 4 °C. The supernatants were then collected, and unbound actin-binding domain fluorescence intensity was analyzed using a fluorimeter (Biotek Instruments, Inc.). Normalized bound fractions were fitted with the following equation 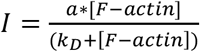. Where *I* is the normalized bound fraction, *a* is the binding stoichiometry, [f-actin] is the actin concentration and *k*_*D*_ is the dissociation constant. For the case of low affinity actin-binding domains, the plectin mutant K278E and utrn-S158C-PEG, *a* was set equal to 1.

### FRET measurements

To investigate structural changes of CH1-CH2 domains we used FRET between GFP fused to the n-terminus of the CH1-CH2 domain-of-interest and alexa 555 maleimide on CH2. S168 was mutated to cysteine for labelling with alexa 555 maleimide (Life Tech) as described above. FRET measurements were made on a fluorescence plate reader in the presence or absence of f-actin (32 μM). FRET was determined at an excitation wavelength of 488nm and an emission wavelength of 575nm. Donor and acceptor bleed-through signals were collected under identical conditions with the GFP-CH1-CH2 domain lacking the acceptor fluorophore and the free alexa 555 dye, respectively. Bleed-through signals were subtracted from the FRET construct fluorescence.

*Single molecule measurements:* Single molecule measurements were made as described previously (Hansen *et al*, 2013; Hayakawa *et al*, 2014). Briefly, we measured the off-rate from binding dwell time histograms using single molecule TIRF microscopy (Fig S4). F-actin filaments were polymerized to a final concentration of 5µM for 1 hour at room temperature and then tethered to pegylated glass surfaces (5% biotin, peg 2k (rapp polymere), (Bieling *et al*, 2010)). Surfaces were assembled in a flow chamber configuration (Bieling *et al*, 2010, 2017) and incubated with streptavidin followed by biotin-phalloidin (Life Technologies) to create a functional surface for tethering actin filaments.. The final buffer for imaging was contained 10µg/mL β-Casein (Sigma) with 0.05nM of binding protein to obtain single molecule dilutions in f-buffer. Images were acquired with TIRF microscopy at a frame rate of 100ms/frame for WT utrn. Due to the slower unbinding kinetics of the mutant Q33A T36A a frame rate of 600ms/frame was used. Single particles were tracked using TrackNTrace (Stein & Thiart, 2016) and analyzed with a custom written MatLab routine.

### Statistics

Error bars represent standard error for relative bound fraction and FRAP measurements. Confidence intervals for fitted data are reported melting temperature measurements. Statistical significance was determined by a two-tailed student’s t-test and assumed significant when p<0.05.

## ACKNOWLEGEMENTS

The authors would like to thank Dyche Mullins and members of the Fletcher lab for helpful discussions. This work was supported by grants from NIH (D.A.F). A.R.H. was in receipt of an EMBO long-term fellowship 1075–2013 and HFSP fellowship LT000712/2014. B.B. was supported by the Ruth L. Kirschstein NRSA fellowship from the NIH (1F32GM115091). P.J was supported by an NSF Fellowship and the Berkeley Fellowship for Graduate Studies. A.B. was supported by the Miller Institute for Basic Research at UC Berkeley as a Miller Visiting Professor. The authors acknowledge the use of the Molecular Imaging Center core facility at UC Berkeley, and the help of the Marqusee Lab with the CD and melting temperature experiments. SAXS data was collected at SIBYLS beamline 12.3.1 at the Advanced Light Source (ALS). SAXS data collection at SIBYLS is funded through DOE BER Integrated Diffraction Analysis Technologies (IDAT) program and NIGMS grant P30 GM124169-01, ALS-ENABLE. D.A.F. is a Chan Zuckerberg Biohub Investigator.

