## Supplementary Figures for "Steric Regulation of Tandem Calponin Homology Domain Actin-Binding Affinity"

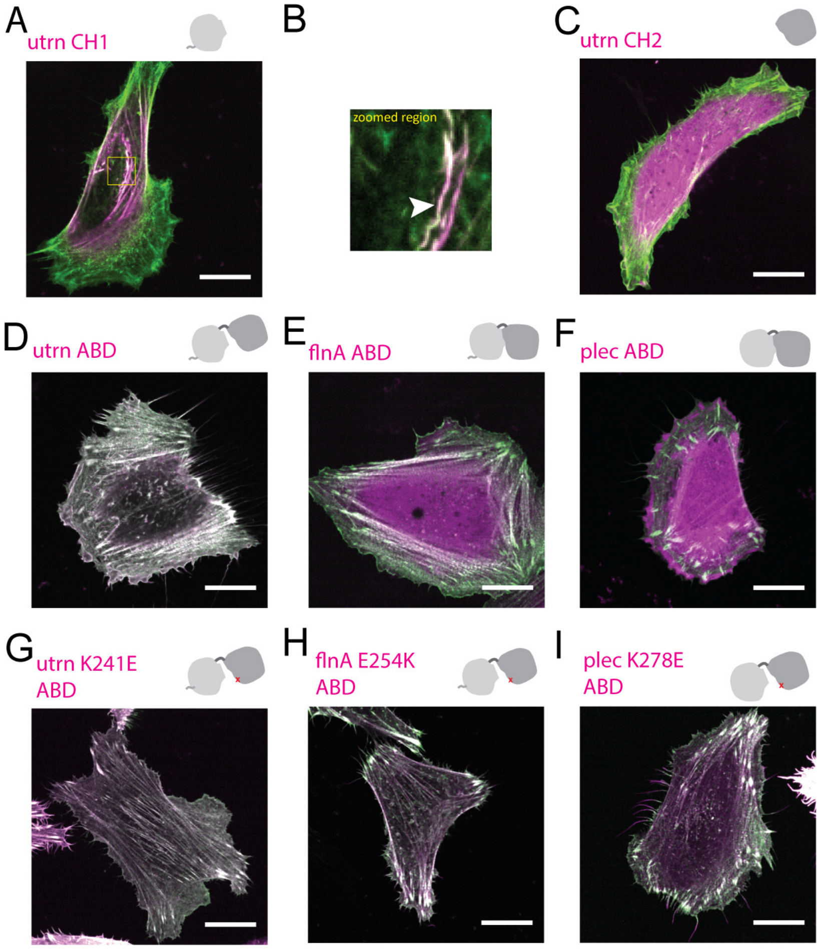


**Figure S1: Images of different CH1-CH2 expressed in live cells relative to the actin-binding domain of utrophin**

In all figures the construct of interest is shown in magenta and the actin-binding domain of utrophin shown in green. Scale bars are 20μm. (A) The CH1 alone from the actin-binding domain of utrophin. (B) Zoomed region with white arrowhead highlighting unstably aggregated protein. (C) The CH2 alone from the actin-binding domain of utrophin. (D) The actin-binding domain of utrophin imaged in both color channels. (E) The CH1-CH2 from filamin A. (F) The CH1-CH2 from plectin. (G-I) Mutations predicted to open the conformation of the CH1-CH2 for utrophin (G, K241E), filamin A (H, E254K) and plectin (I, K278E).

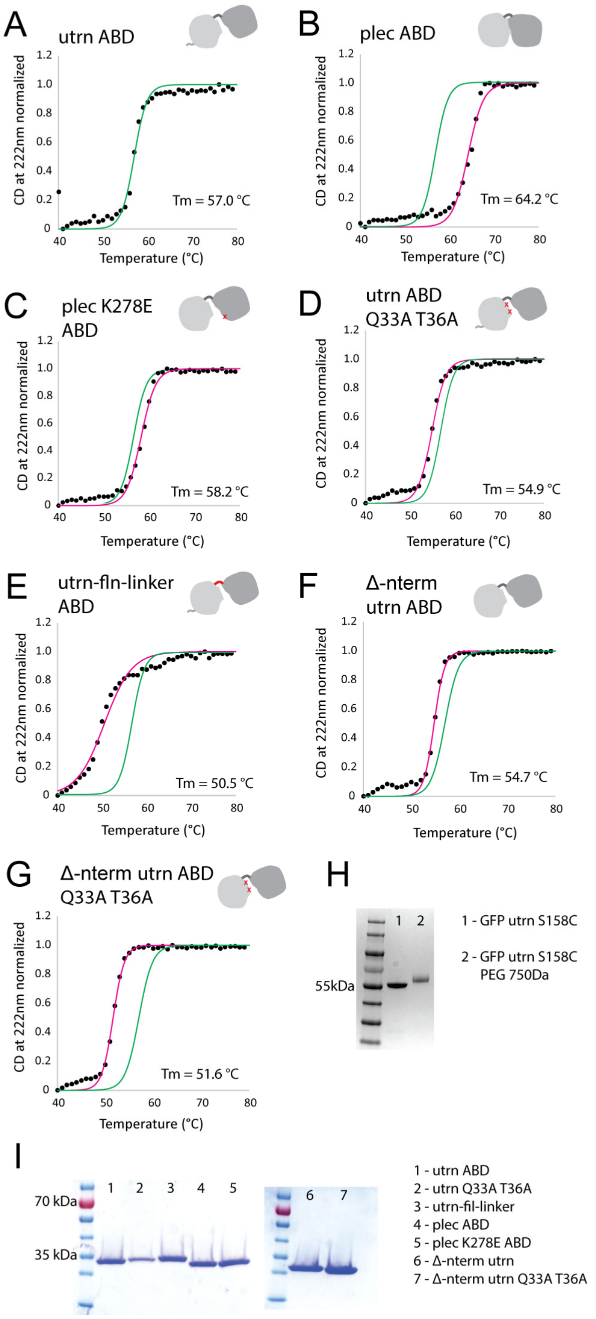


**Figure S2: *In vitro* characterization of actin-binding domain properties**

In all plots, measured values shown as black points and fitted curves as solid lines. The fitted curve for utrophin actin-binding domain is shown in all plots in green as a reference and CH1-CH2 of interest shown in magenta. Melting curves for the actin-binding domain of (A) utrn, (B) plectin, (C) plectin mutant, (D) utrn Q33A T36A, (E) utrophin-fil-linker, (F) Δ-nterm, (G) Δ-nterm Q33A T36A. (H) SDS page for PEG conjugated actin binding domains. (I) SDS page for actin binding domains used in this study.

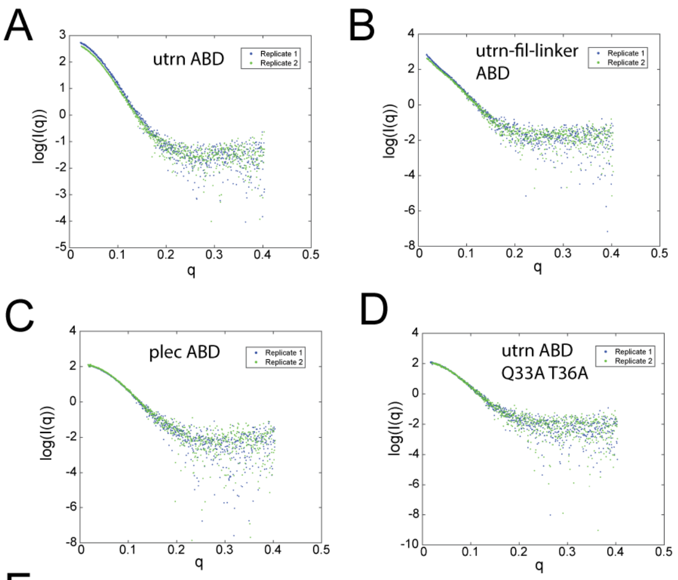


E

| Sample | Buffer | Rc | Rg_inv | Rg_real | dmax | P_x | MW |
|---|---|---|---|---|---|---|---|
| utrn ABD_a | 1 | 16.1 | 23.5 | 23.26 | 100.5 | 3.4 | 2.70E+04 |
| | 2 | 16.1 | 24.1 | 23.89 | 111 | 3.6 | 2.90E+04 |
| utrn ABD_b | 1 | 16.1 | 23.5 | 23.47 | 86.5 | 3.3 | 2.60E+04 |
| | 2 | 16.2 | 23.7 | 23.69 | 87.5 | 3.4 | 2.70E+04 |
| utrn-fil-linker ABD_a | 1 | 16.4 | 42 | 41.92 | 146 | 3.1 | 4.40E+04 |
| | 2 | 16.6 | 44.8 | 44.88 | 154 | 3.8 | 4.40E+04 |
| utrn-fil-linker ABD_b | 1 | 16.9 | 38.8 | 38.74 | 133.5 | 2.9 | 3.70E+04 |
| | 2 | 16.3 | 39.1 | 39.08 | 136 | 3.2 | 3.80E+04 |
| plect ABD_a | 1 | 15.9 | 21.9 | 21.86 | 85.5 | 3.5 | 2.30E+04 |
| | 2 | 15.4 | 21.9 | 21.79 | 81.5 | 3.4 | 2.70E+04 |
| plect ABD_b | 1 | 15.6 | 21.6 | 21.53 | 76 | 3.2 | 2.30E+04 |
| | 2 | 15.8 | 21.6 | 21.57 | 72.5 | 3.3 | 2.80E+04 |
| utrn Q33A T36A ABD_a | 1 | 15 | 23.4 | 23.38 | 82 | 3 | 2.60E+04 |
| | 2 | 15.8 | 23.5 | 23.43 | 80.5 | 3.3 | 3.00E+06 |
| utrn Q33A T36A ABD_b | 1 | 15.9 | 23.4 | 23.41 | 78 | 3.1 | 2.50E+04 |
| | 2 | 15.8 | 23.6 | 23.55 | 81 | 3.4 | 3.10E+04 |

**Figure S3: Small Angle X-ray Scattering (SAXS) data**

(A-D) SAXS intensity plots are shown for two replicates of each sample. The replicates are in high agreement with each other. The overall profiles indicate that utrophin-filamin-linker ABD is qualitatively different in total shape from utrn ABD, plec ABD, and utrn ABD Q33A T36A. (E) Guinier analysis, P(r) distribution estimation, and Porod volume and exponent estimations were performed to determine cross-sectional radius of gyration (Rc), radius of gyration (from Guinier, Rg_inv, from P(r), Rg_real), dmax, Porod exponent (P_x), and molecular weight (MW). Rg_real values indicate that utrophin-filamin-linker ABD is larger, and plectin ABD is smaller, than WT utrophin ABD. WT utrophin ABD and Q33A T36A ABD appear similar in overall size.

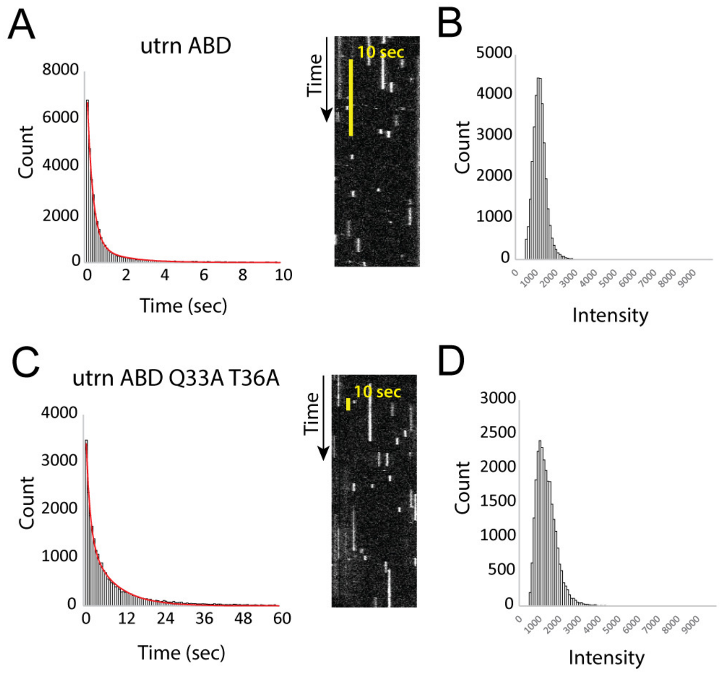


**Figure S4: Single molecule binding measurements**

(A) Binding dwell time histogram for utrn ABD and kymograph (right). Scale bar is 10 seconds. (B) Binding intensity histogram for utrn ABD. (C) Binding dwell time histogram for utrn Q33A T36A and kymograph (right). Scale bar is 10 seconds. (D) Binding intensity histogram for utrn Q33A T36A.

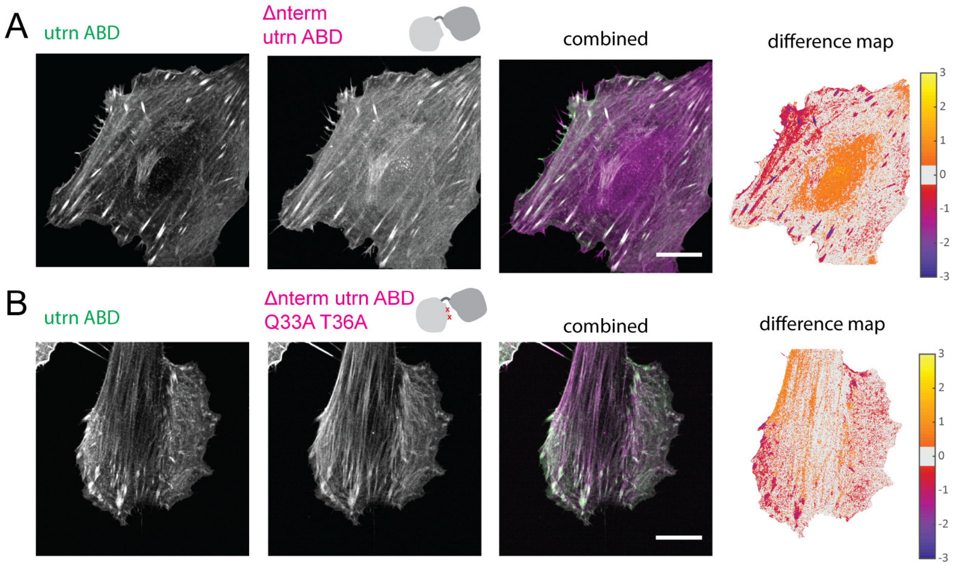


**Figure S5: Loss in binding through n-terminal truncation recovered by introducing CH1-CH2 interface mutations**

(A) N-terminal flanking region truncation construct from utrophin. (B) N-terminal flanking region truncation construct from utrophin plus the mutations Q33A T36A. Scale bars are 20μm.

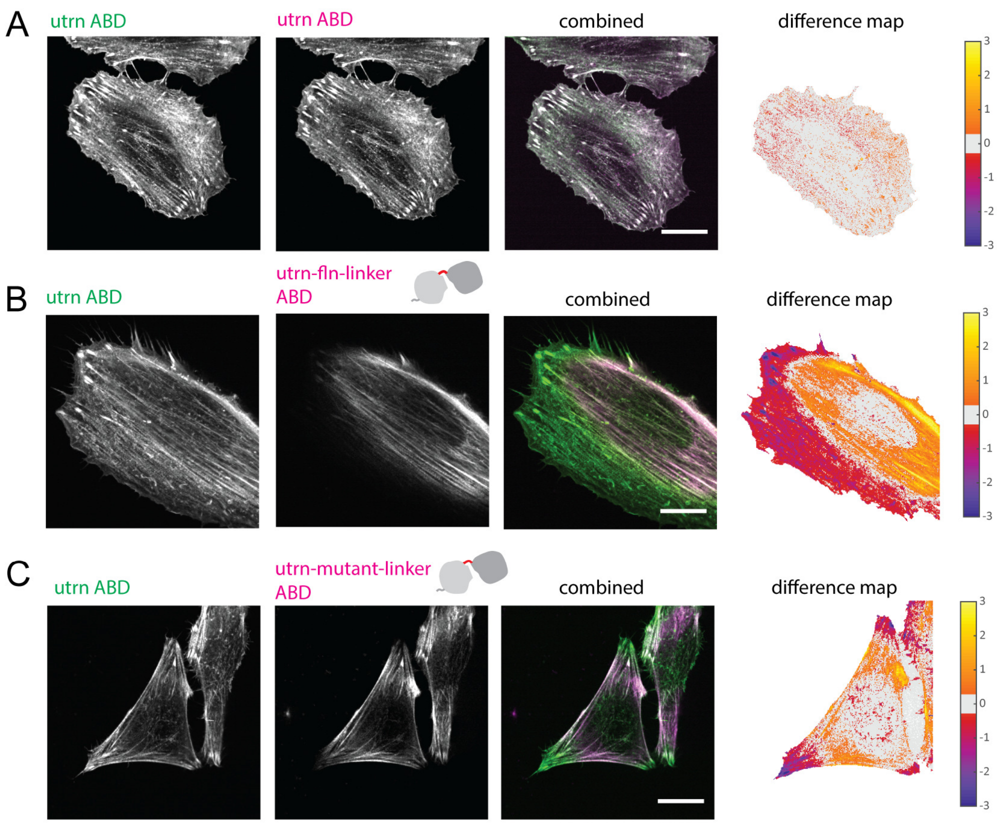


**Figure S6: Images of different CH1-CH2 in live cells**

(A) The actin-binding domain of utrophin in both color channels, (B) utrophin-filamin-linker shown in magenta, (C) utrophin with mutated linker region shown in magenta. Difference maps are shown to the right. Scale bars are 20μm.

A

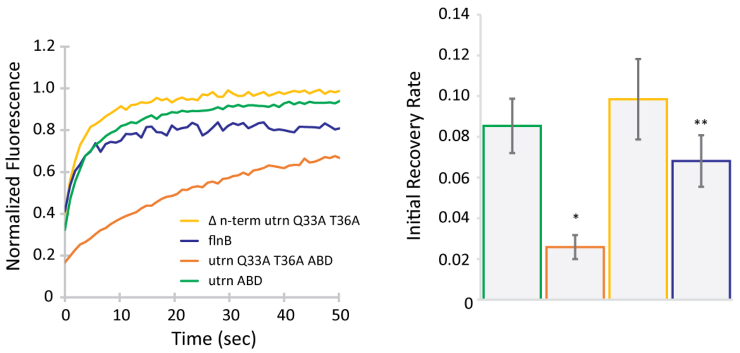


**Figure S7: Dynamics of different CH1-CH2 measured by Fluorescence Recovery After Photobleaching (FRAP)**

(A) Fluorescence recovery curves for different CH1-CH2 constructs (left) and the initial rates of recovery measured from these curves as an estimate of the dynamic exchange of proteins on actin (right) for WT utrophin ABD (green), filamin B (blue, p**<0.05), Q33A T36A (orange, p*<0.05) and Δ-nterm Q33A T36A (yellow, p=0.05).

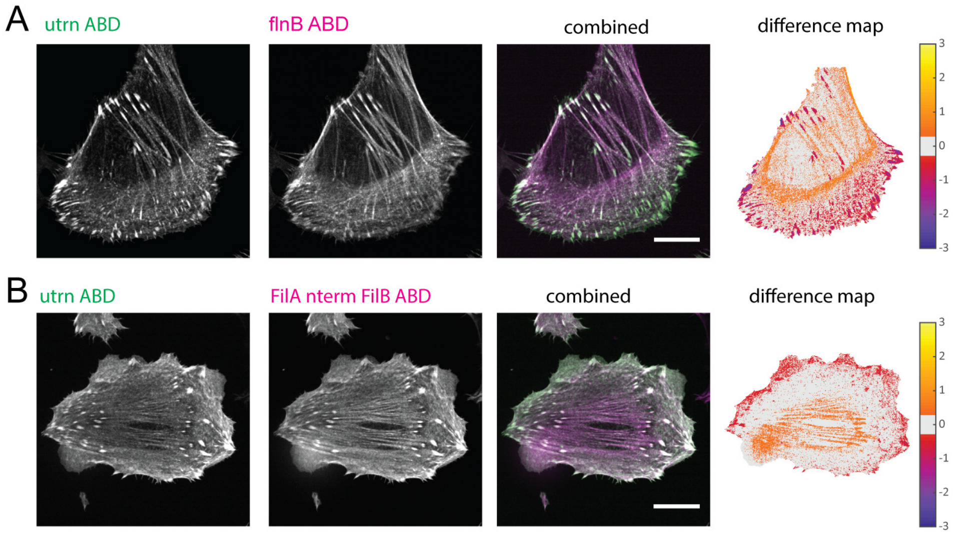


**Figure S8: Filamin B and n-terminal chimera with filamin A**

(A) CH1-CH2 from filamin B (magenta). (E) A chimera containing the N-terminal flanking region from filamin A and the CH1-CH2 from filamin B (magenta). Scale bars are 20µm.
